# C-terminal binding proteins 1 and 2 in traumatic brain injury-induced inflammation and their inhibition as an approach for anti-inflammatory treatment

**DOI:** 10.1101/820258

**Authors:** Hong Li, Caiguo Zhang, Chunxia Yang, Melanie Blevins, David Norris, Rui Zhao, Mingxia Huang

## Abstract

Traumatic brain injury (TBI) induces an acute inflammatory response in the central nervous system that involves both resident and peripheral immune cells. The ensuing chronic neuroinflammation causes cell death and tissue damage and may contribute to neurodegeneration. The molecular mechanisms involved in the maintenance of this chronic inflammation state remain underexplored. C-terminal binding protein (CtBP) 1 and 2 are transcriptional coregulators that repress diverse cellular processes. Unexpectedly, we find that the CtBPs can transactivate a common set of proinflammatory genes both in lipopolysaccharide-activated microglia, astrocytes and macrophages, and in a mouse model of the mild form of TBI. We also find that the expression of these genes is markedly enhanced by a single mild injury in both brain and peripheral blood leukocytes in a severity- and time-dependent manner. Moreover, we were able to demonstrate that specific inhibitors of the CtBPs effectively suppress the expression of the CtBP target genes and thus improve neurological outcome in mice receiving single and repeated mild TBIs. This discovery suggests new avenues for therapeutic modulation of the inflammatory response to brain injury.

## Introduction

Traumatic brain injury (TBI) is a leading cause of death and disability among children and young adults in the United States (1–3). It can be classified as mild, moderate and severe based on neurological injury severity. Severe TBI is a well-established risk factor for several diseases of the aging brain, including amyotrophic lateral sclerosis, Alzheimer’s disease and Parkinson’s disease (4). The mild form of traumatic brain injury (mTBI, also known as concussion) accounts for >80% of all TBIs (5, 6). The majority of mTBI patients experience full recovery within a couple of weeks to months. A small minority of these patients (10–20%) experience persistent postinjury symptoms (7–9). Disturbingly, even a single mTBI induces pathophysiological changes in the brain that can be detected in both acute and chronic phases postinjury (10–12). Athletes and military service members are especially vulnerable to multiple mTBIs, raising concerns about cumulative or long-term effects of recurrent insults (8, 13). In effect, repeated mTBI has been linked to late-onset neurodegenerative disease development, such as chronic traumatic encephalopathy (14, 15). There are currently no FDA-approved drugs specifically for TBI.

TBI elicits a distinct inflammatory response in the central nervous system (CNS) that involves both local and peripheral immunocytes (16–18). The acute phase of the injury begins within minutes of the primary mechanical damage to the brain tissue. Microglial and astroglial activation can persist for months and years after TBI and can affect CNS neurodegeneration. In mTBI, diffuse axonal injury, often resulting from impact/acceleration forces, leads to the release of danger signals such as damage-associated molecular patterns and alarmins from the stressed axons (19, 20). These signals induce the rapid activation of resident microglia, which upregulate the expression of an array of inflammatory mediators to amplify the immune response and recruit peripheral immune cells to the damage site. Excessive inflammation causes cell death and tissue damage and in the chronic phase may contribute to neurodegeneration (21). There is a compelling need for effective neuroprotective interventions that improve functional outcomes in TBI patients (22, 23). In spite of its recognized importance in the regulation of inflammatory responses, inhibition of NF-κB showed a limited neuroprotective effect in experimental models of TBI (24, 25). As such, the discovery and development of new modulators of brain injury-mediated neuroinflammatory response are the subject of intense pharmaceutical efforts.

C-terminal binding protein (CtBP) 1 and 2 are paralogous transcriptional coregulators that repress diverse biological processes by directly binding transcription factors and recruiting chromatin-modifying enzymes to target gene promoters (26, 27). These two proteins exhibit a remarkable structural similarity to D2-hydroxyacid dehydrogenases, including an N-terminal substrate binding domain, a central NAD(H)-binding domain and a C-terminal extension (28). They also contain a hydrophobic cleft within the N-terminal substrate binding domain known as the PXDLS-binding site and a surface groove within the central NAD(H)-binding domain that recognizes the sequence RRTGXPPXL (RRT motif) (29–31). These two unique structural elements constitute the majority of CtBP interactions with transcription factors and chromatin modifying enzymes such as histone deacetylases, acetyltransferase, methyltransferases and demethylases (32–35). NAD(H) facilitates the dimerization of the CtBPs and promotes their transcriptional corepression function (36, 37). The CtBPs, which were originally identified through their binding to the adenovirus E1A oncoprotein (38), have been demonstrated to regulate multiple transcription factor networks with roles in tumorigenesis and cell survival (26, 39, 40). Inhibition of the interaction between the CtBPs and their PXDLS-containing target proteins by the peptide Pep1-E1A has been shown to block transcriptional repressor activity and mitigate oncogenic phenotypes in mice (41). On the other hand, the small molecule MTOB that inhibits the dehydrogenase activity of the CtBPs attenuates the growth and self-renewal of cancer stem cells in vitro and decreases tumor burden in a mouse model of colon cancer (42–44). In addition, it is postulated that the CtBPs also function as context-dependent transcriptional coactivators (34, 45, 46). In this report, our results indicate that CtBP1 and CtBP2 increase the in vivo transcription of proinflammatory genes in response to mTBI; reveal that the mechanism underlying both local and systematic induction of these CtBP target genes operates in an impact energy dose- and time-dependent manner; and indicate that Pep1-E1A and MTOB inhibit the transactivating activity of the CtBPs, reduce microglia activation and ameliorate neurological deficits following single and repeated injuries in mice, thus demonstrating the potential utility of these agents to modulate the immune response to head injury.

## Results and Discussion

### CtBP1 and CtBP2 are required for the transactivation of proinflammatory genes in LPS-activated microglia and macrophages

To explore the role of the CtBPs in cellular regulation of the inflammatory response, we investigated their occupancy on the promoters of the 674 inflammatory response genes of the Gene Ontology GO:0006954 set (47, 48) through the analysis of CtBP ChIP-seq data obtained from non-stimulated breast cancer cells (49). We observed a moderate enrichment in the promoters of the *IL1B*, *IL6*, *TNFA, ICAM1, VCAM1, S100A8, S100A9, NLRP3* and *PTGS2* genes. To assess whether the inflammation-induced expression of the above genes is CtBP-dependent, we evaluated the effects of siRNA-mediated simultaneous knockdown of *CtBP1* and *CtBP2* on the mRNA levels of these candidate genes in lipopolysaccharide (LPS)-stimulated microglia and macrophage using the reverse transcription and quantitative real-time PCR (RT-qPCR) method. As shown in Figures 1A and 1B, exposure to LPS induces the expression of all nine genes (3- to 70-fold increase) in the murine BV2 microglia and RAW264.7 macrophages. Silencing of the CtBP genes markedly decreases the LPS-induced gene expression, ranging from 40% to 85% (see Figures 1A and 1B), suggesting that CtBP1 and CtBP2 are involved in the upregulation of these genes in response to LPS activation. We also observed similar CtBP-dependent activation of the above nine genes in LPS-stimulated human THP-1 monocytes. Thus, CtBPs likely regulate the expression of the nine candidate genes in the human and mouse monocytes. To validate our analysis, we performed chromatin immunoprecipitation (ChIP) experiments in the LPS-activated BV2 cells to measure CtBP binding to the promoter regions of the four strongly induced genes (*IL1B, IL6, TNFA* and *S100A8*) as a testbed. Indeed, LPS stimulates a significant increase in the binding of CtBP1 (10- to 17-fold change) and CtBP2 (7- to 14-fold change) to the promoter regions of these genes (Figure 1C). We also found that the promoter recruitment of histone acetyltransferase (HAT) p300, a PXDLS-containing protein that binds CtBP (32), is intimately associated with CtBP occupancy at the four target genes (Figure 1C). Taken together, our findings indicate that CtBP1 and CtBP2 serve as transcriptional coactivators capable of enhancing LPS-induced expression of the nine aforementioned proinflammatory genes.

**Figure 1.**
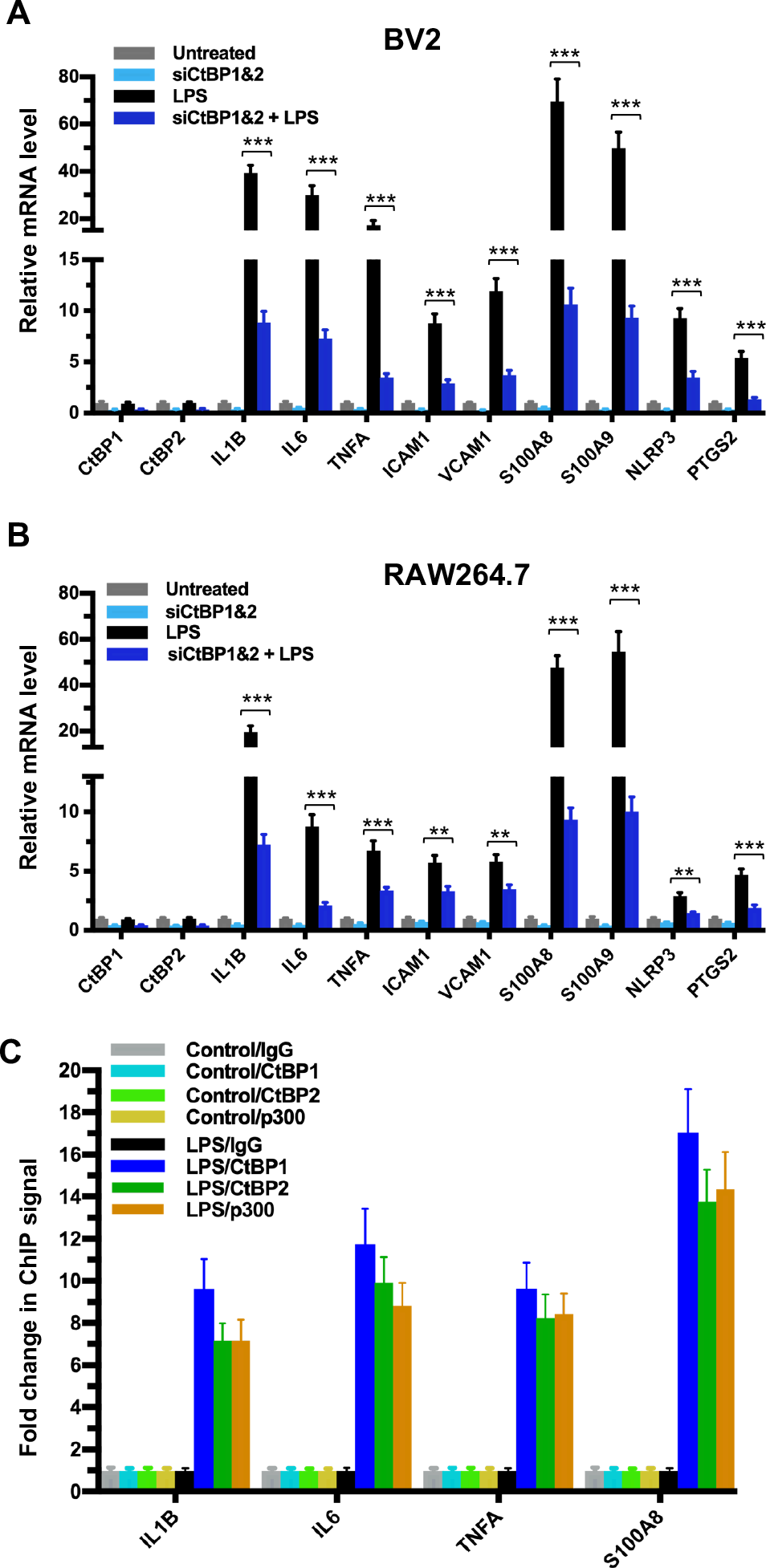
CtBP1 and CtBP2 are required for LPS-induced proinflammatory gene expression in microglia and macrophage cell lines. (**A–B**) Simultaneous knockdown of CtBP1 and CtBP2 suppresses mRNA expression of proinflammatory genes in LPS-activated microglia and macrophages. Murine RAW264.7 (**A**) and BV2 (**B**) cells were transfected with siRNAs specific for CtBP1 and CtBP2 or scrambled control siRNAs for a period of 24 h, followed by LPS stimulation (200 ng/ml) for 6 h. Total RNA was extracted and analyzed by the RT-qPCR. Relative mRNA expression was normalized to that of *ACTB* and depicted as fold changes vs. scrambled siRNA transfected and non-stimulated control (Untreated). Data are presented as mean ± SD. *n* = 3; \*\**p*<0.01, \*\*\**p*<0.001. (**C**) LPS induces increased binding of CtBP1, CtBP2 and p300 to the *IL1B, IL6, TNFA* and *S100A8* gene promoters. Chromatin fractions from control and LPS-treated BV2 cells were precipitated with antibodies specific to CtBP1, CtBP2 and p300. Bars represent fold changes of relative ChIP signals normalized to the respective controls.

### Pep1-E1A suppresses the induction of the CtBP target genes in LPS-activated mouse primary microglia and astrocytes

We have shown previously that the Pep1-E1A peptide disrupts the interaction of CtBP1 with the PLDLS-containing repressor ZEB1 in cancer cells and relieves the transcriptional repression of *CDH1* (E-cadherin) and *BAX* by the CtBPs (Blevins et al., 2018). To explore whether this peptide inhibitor can interfere with the transactivation function of CtBP1 and CtBP2, we preincubated Pep1-E1A with mouse primary microglia and astrocytes for 2 h prior to stimulation with LPS and measured mRNA levels of the above nine CtBP-regulated proinflammatory genes by RT-qPCR. As expected, the LPS treatment of both microglia and astrocytes increases the expression of these nine genes by 7- to 132-fold while reducing the expression of *CDH1* and *BAX* to approximately 50% of the levels seen in controls (Figures 2A and S1A). In contrast, Pep1-E1A pretreatment significantly suppresses the LPS-induced expression of the nine proinflammatory genes by 40 to 70%, and effectively relieves the repression of *CDH1* and *BAX* by ~50% (Figures 2A and S1A). We also note that Pep1-E1A pretreatment results in a 40% to 50% decreases in the basal mRNA expression of these proinflammatory genes in both primary microglia and astrocytes without stimulation with LPS, and causes a 57% and 80% increase in the basal mRNA levels of *CDH1* and *BAX*, respectively (Figures 2A and S1A). We conclude that CtBP1 and CtBP2 play a crucial role in mediating the activation of the nine proinflammatory genes. These findings also indicate that the CtBPs exhibit a dual role in transcriptional activation and repression.

**Figure 2.**
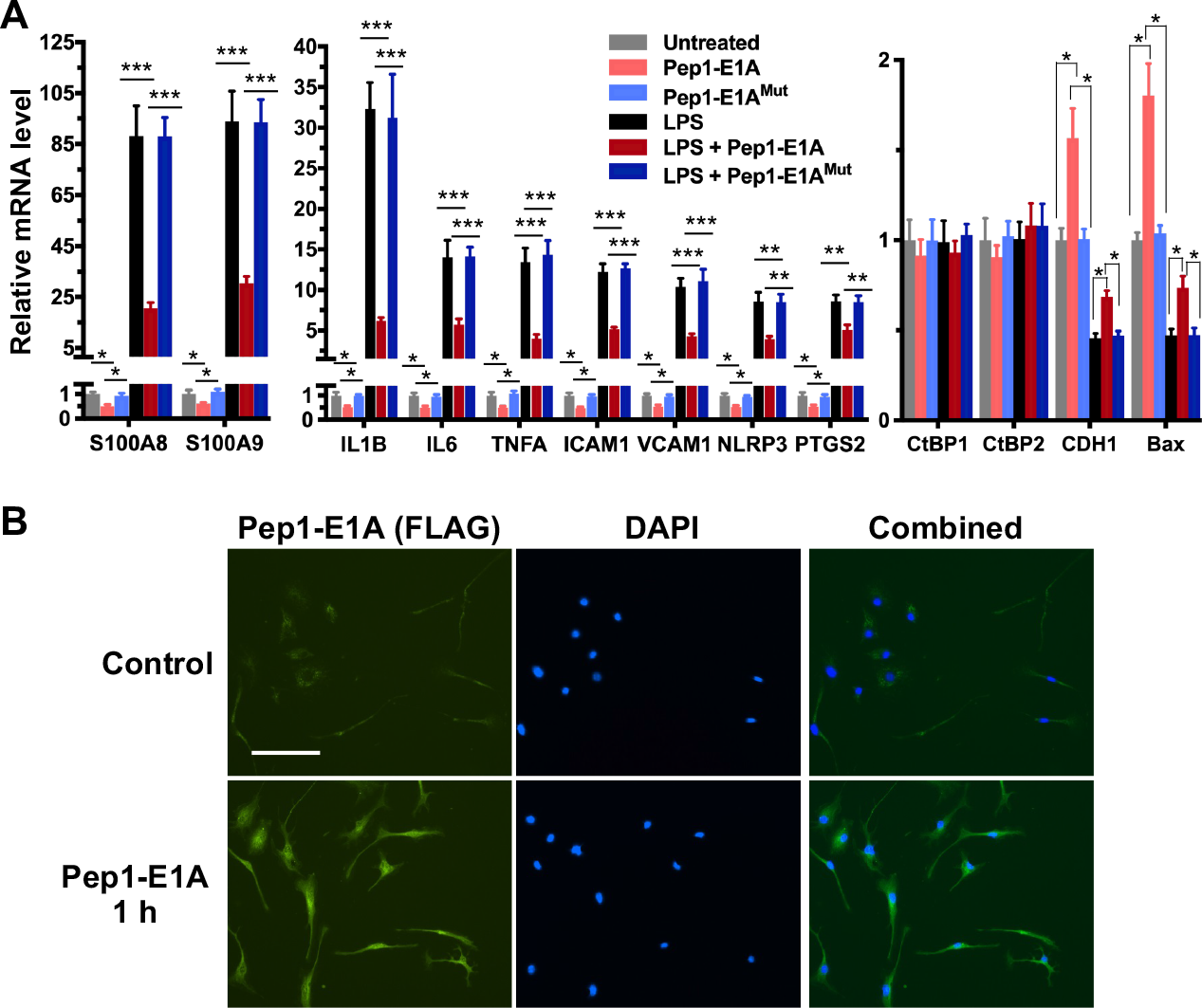
Pep1-E1A suppresses the induction of CtBP-regulated proinflammatory genes in LPS-activated mouse primary microglia. (**A**) Mouse primary microglia were incubated with 20 μM Pep1-E1A or the PLDLS-to-PLDEL mutant (Pep1-E1A^Mut^) peptides for 2 h, followed by LPS stimulation (200 ng/ml) for 2 h. Relative mRNA expression was depicted as fold changes vs. the untreated controls. *n* = 3; \**p*<0.05, \*\**p*<0.01, \*\*\**p*<0.001. (**B**) Representative immunostaining images of Pep1-E1A internalization into cultured primary microglia after 1 h incubation with the peptide. Scale bar, 50 μm.

### Mild TBI causes dynamic changes in the induction of gene expression by CtBP1 and CtBP2 in both brain and peripheral blood leukocytes

To directly evaluate the hypothesis that the CtBPs play a role in the regulation of the acute inflammatory response following mild TBI, we used a closed-head impact model of engineered rotational acceleration (CHIMERA) in mice to investigate the dose–response relationship between injury severity and changes in the expression of CtBP target genes in vivo. This mouse model was selected in the present study because diffuse axonal injury is the hallmark of inertial TBI (50). Animals received sham or a single head injury with the impact kinetic energy of 0.5 J, 0.65 J and 0.8 J, respectively. At 24 h postinjury, brain and blood leukocyte samples were collected to measure mRNA levels of the five CtBP-regulated proinflammatory genes identified above as well as those of *CtBP1* and *CtBP2* by RT-qPCR. The single brain injury leads to the increased expression of all five proinflammatory genes and *CtBP2*, in both CNS and systemic compartments, in an impact energy dose-dependent manner (Figures 3A and 3B). Specifically, the *NLRP3*, *ICAM1*, *PTGS2* and *CtBP2* genes are moderately upregulated in the mouse brain, with 2.2- to 2.9-fold change at 0.5 J, 3.2- to 3.7-fold change at 0.65 J and 4.0- to 4.4- fold change at 0.8 J. The *IL1B* gene is induced to a significant degree, with 23-, 34-, and 45-fold change at 0.5 J, 0.65 J and 0.8 J, respectively. The expression of *S100A8* is exceptionally high (76-, 93- and 109-fold change at 0.5 J, 0.65 J and 0.8 J, respectively). Remarkably, there is a strong, linear correlation between the mRNA expression levels of the above six genes in the mouse brain and peripheral blood leukocyte samples (Figures 3A and 3B). In parallel, we observed impact energy dose-dependent increases in the expression of the S100A8, NLRP3 and CtBP2 proteins in the injured mouse brain, as assessed by the Western blot method (Figure 3C). These data demonstrate the dose-dependent effects of mild brain injury on CtBP-mediated inflammatory response in mice. In addition, the upregulated *CtBP2* gene expression observed following mTBI has implications for the mechanism of CtBP-mediated transcriptional activation (see Figures 3 A–C).

**Figure 3.**
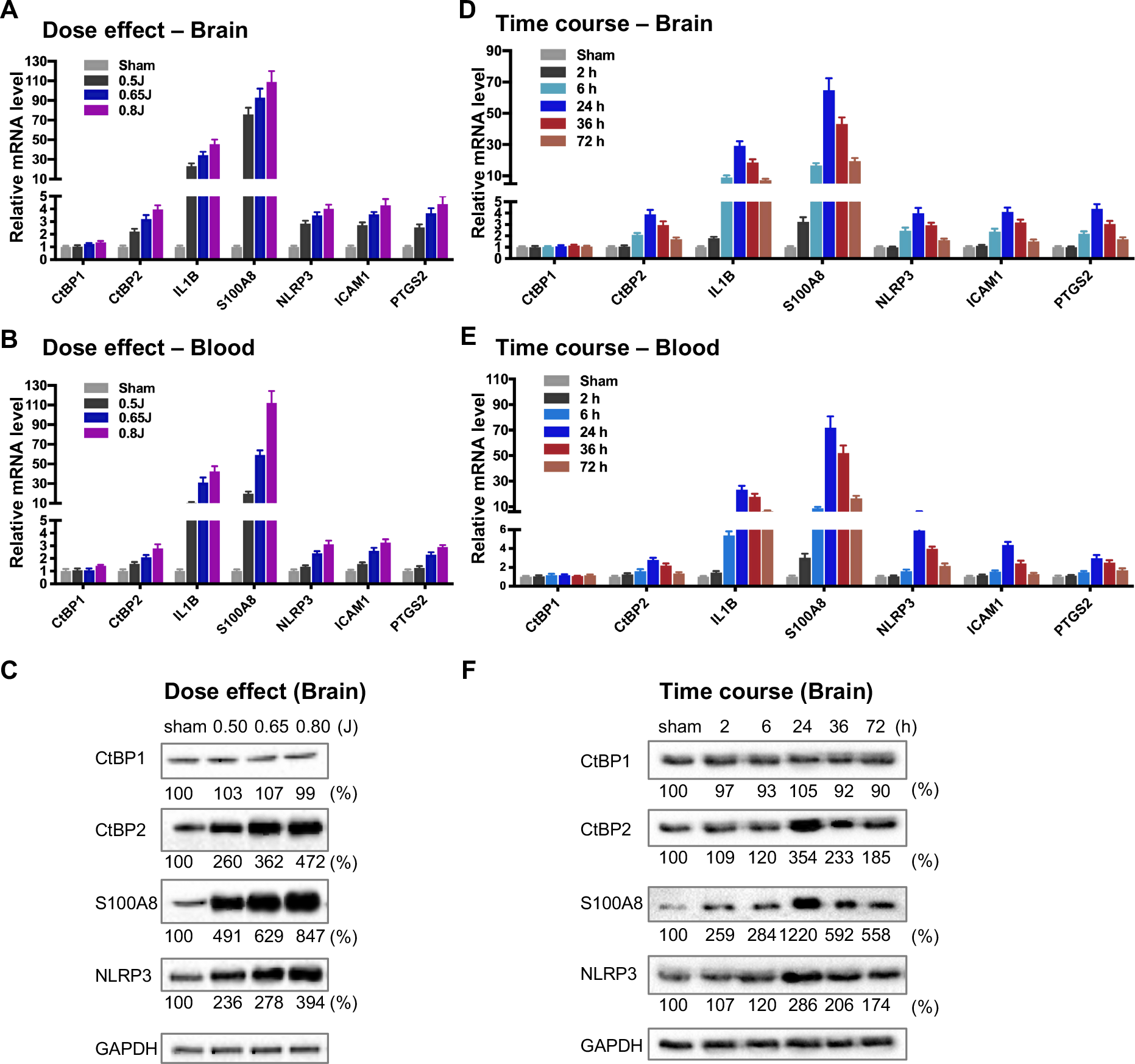
Impact dose- and time-dependent induction of CtBP target genes in brain and peripheral blood leukocytes following single mild TBI. (**A**and **B**) Mice (*n* = 5 per group) were subjected to a single head injury of varying impact energies and the brain (**A**) and peripheral blood leukocytes (**B**) were harvested for mRNA analysis 24 h after injury. Results were normalized to the sham group and are presented as mean ± SD. (**C**) Representative Western blots of the CtBP1, CtBP2, S100A9, NLRP3 proteins of the brain tissues from the dose-response groups. Relative protein expression was normalized to the loading control GAPDH and shown as percent change vs sham under the blots. (**D**and **E**) Mice (*n* = 5 per group) received a single head impact of 0.7 J energy and mRNA expression in brain (**D**) and blood (**E**) were analyzed at the indicated time points postinjury. (**F**) Western blots showing protein expression of the brain tissues from the time course experiment.

To characterize the kinetics of the above head injury-induced gene expression, animals received a single impact of 0.7 J energy and were euthanized at different time points afterwards. The mRNA expression levels of the five proinflammatory genes and *CtBP2* started to increase in both brain and peripheral blood leukocytes as early as 2 h postinjury, peaked at 24 h, and then gradually decreased, but the level remained significantly higher than that in the sham control 72 h after injury (*p* < 0.05, Figures 3D and 3E). Western blot analysis also shows time-dependent changes in the expression of the S100A8, NLRP3 and CtBP2 proteins in the injured brain (Figure 3F). Moreover, the highest levels of these protein expression coincide with those of their mRNAs at 24 h postinjury (see Figures 3D and 3F). Our data suggest that the CtBPs may play a role in determining the timing and intensity of the inflammatory response to brain injury, consistent with cerebral and plasma proinflammatory cytokine profile following injury in human TBI patients (51, 52).

To address the question whether the TBI-induced proinflammatory gene expression observed in the peripheral blood leukocytes (see Figures 3B and 3E) is caused by the release of inflammatory mediators from the injured brain or the region of the skin and/or skull at the point of impact, three groups of mice received single head, back and ear impacts at 0.7 J energy, and their blood samples and skin tissues containing the site of the impact were collected 24 h postinjury. Moderate mRNA expression levels of the nine CtBP target genes identified in the present study were observed in the skin of the three injury groups, whereas the head injury group exhibits significantly increased mRNA levels of these genes in the circulating leukocytes relative to the other two groups (*p* < 0.001, Figures S2A and S2B). In particular, the most strongly upregulated genes *S100A8* and *S100A9* in circulating leukocytes display >95-fold change in the head injury group, and 6- to 8- fold change in the back and ear injury groups (Figure S2B). Our results suggest that the single mild head injury can evoke a systemic inflammatory response. Consistent with this argument, systemic inflammatory response syndrome is frequently observed in human patients with TBI (53). We therefore propose that CtBP1 and CtBP2 are potential mediators of both local and systemic inflammation in response to mTBI.

### Therapeutic benefits of CtBP inhibition after mTBI

Because Pep1-E1A suppresses LPS-induced inflammatory response in primary microglia and astrocytes by directly targeting the transcriptional activation function of CtBP1 and CtBP2 (see Figures 2A and S1A), we investigated whether Pep1-E1A could be useful in attenuating the acute inflammatory sequelae of brain trauma using our mouse model of mild head injury. To assess the therapeutic efficacy of Pep1-E1A on head trauma, animals were subjected to the application of a single head impact of 0.8 J energy. The animals were then randomized to begin treatment with placebo (vehicle), Pep1-E1A or Pep1-E1A^Mut^ (3 mg/kg by i.p. injection at 1 h and 24 h postinjury). An evaluation of neurological function was performed at 1 h, 24 h and 48 h after injury using criteria of the neurological severity score (NSS). We found that averaged NSS numbers were statistically different between the vehicle control and Pep1-E1A-treated groups at 48 h postinjury (*p* < 0.05), while no significant differences were observed between the vehicle control and Pep1-E1A^Mut^-treated groups (Figure 4A). As expected, Pep-E1A treatment significantly reduces mRNA expression of the nine CtBP-regulated proinflammatory genes described above near 48 h postinjury in both brain tissue and peripheral blood leukocytes (Figures 4A and 4B). Thus, these results demonstrate that the inhibition of CtBP1 and CtBP2 ameliorates mTBI-induced inflammatory damage in mice. It is also noteworthy that the inhibitory effect of Pep1-E1A on the CtBP-mediated transactivation is more pronounced in the peripheral blood cells (56–83%) than in the brain tissues (21–41%), presumably due to the inability of the peptidic Pep1-E1A inhibitor to efficiently cross the blood-brain barrier.

**Figure 4.**
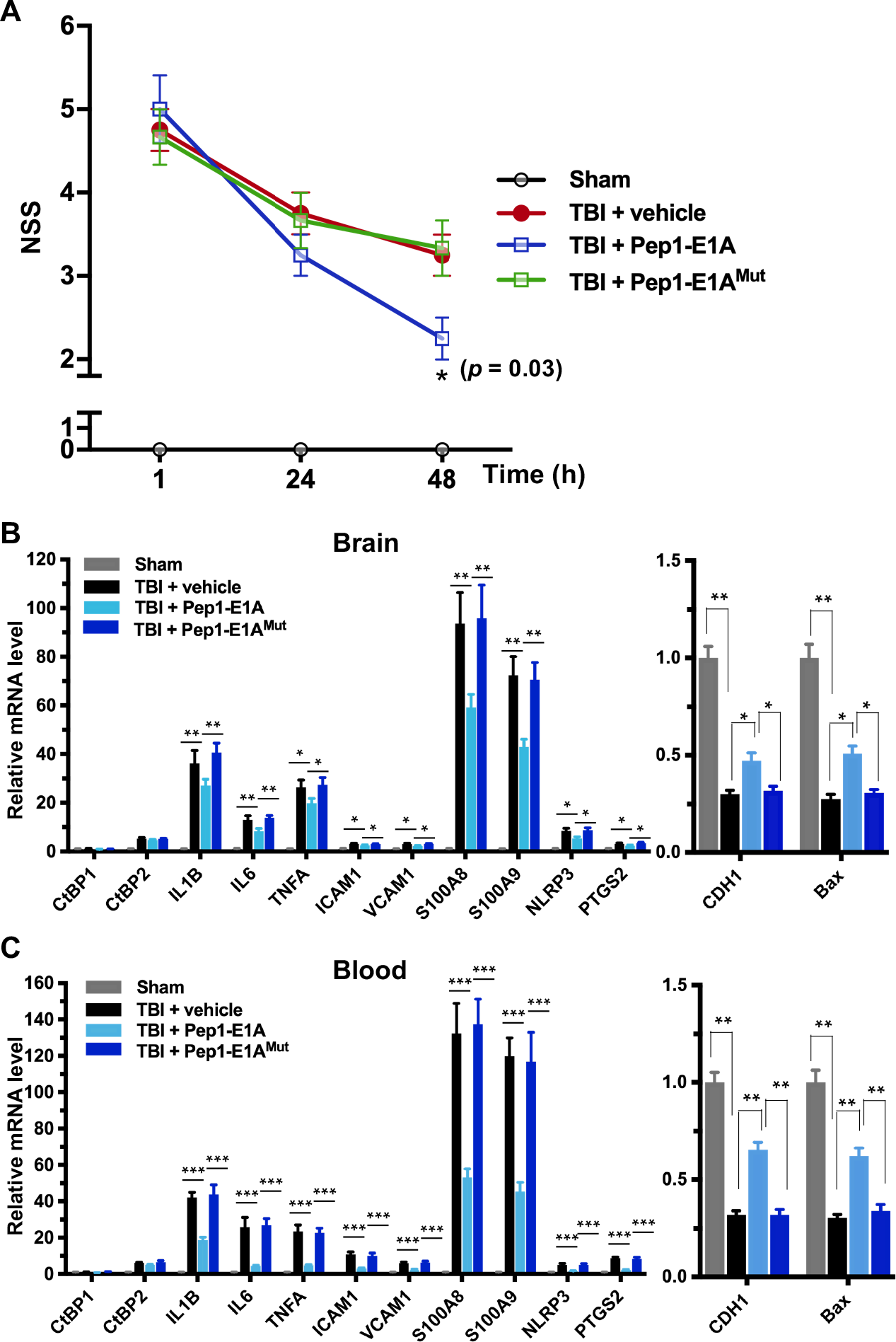
Pep1-E1A reduces CtBP-dependent proinflammatory gene expression and ameliorates neurological deficits after mild brain injury. (**A**) Comparison of NSS scores at 1, 24 and 48 h postinjury. *n* = 4 or 5; \**p*<0.05. (**B**and **C**) Comparison of mRNA expression of the CtBP target genes in brain (**B**) and peripheral blood leukocytes (**C**) at 48 h postinjury in mice that received sham, a single 0.8 J TBI, or TBI followed by treatment with Pep1-E1A or Pep1-E1A^Mut^ (2 mg/kg). Results (mean ± SD) were normalized to sham. *n* = 4 or 5; \**p*<0.05, \*\**p*<0.01, \*\*\**p*<0.001.

The small molecule NSC95397, like the peptide Pep1-E1A, inhibits the transcriptional repressor activity of CtBP1 and CtBP2 by interfering with their binding to PXDLS-containing targets (Blevins et al., 2015). The above mouse model of mild head injury was used to examine the effectiveness of NSC95397 in limiting neuronal damage following TBI. NSC95397-treated mice showed a significant improvement in neurobehavioral deficits, and lower NSS compared to the vehicle control at 48 h (*p* < 0.05) and 72 h (*p* < 0.01) postinjury (Figure 5A). NSC95397 demonstrated equal inhibitory effects on the expression of the nine CtBP target genes in brain tissue (66–86%) and peripheral blood leukocytes (68–90%) at 72 h postinjury (Figures 5B and 5C).

**Figure 5.**
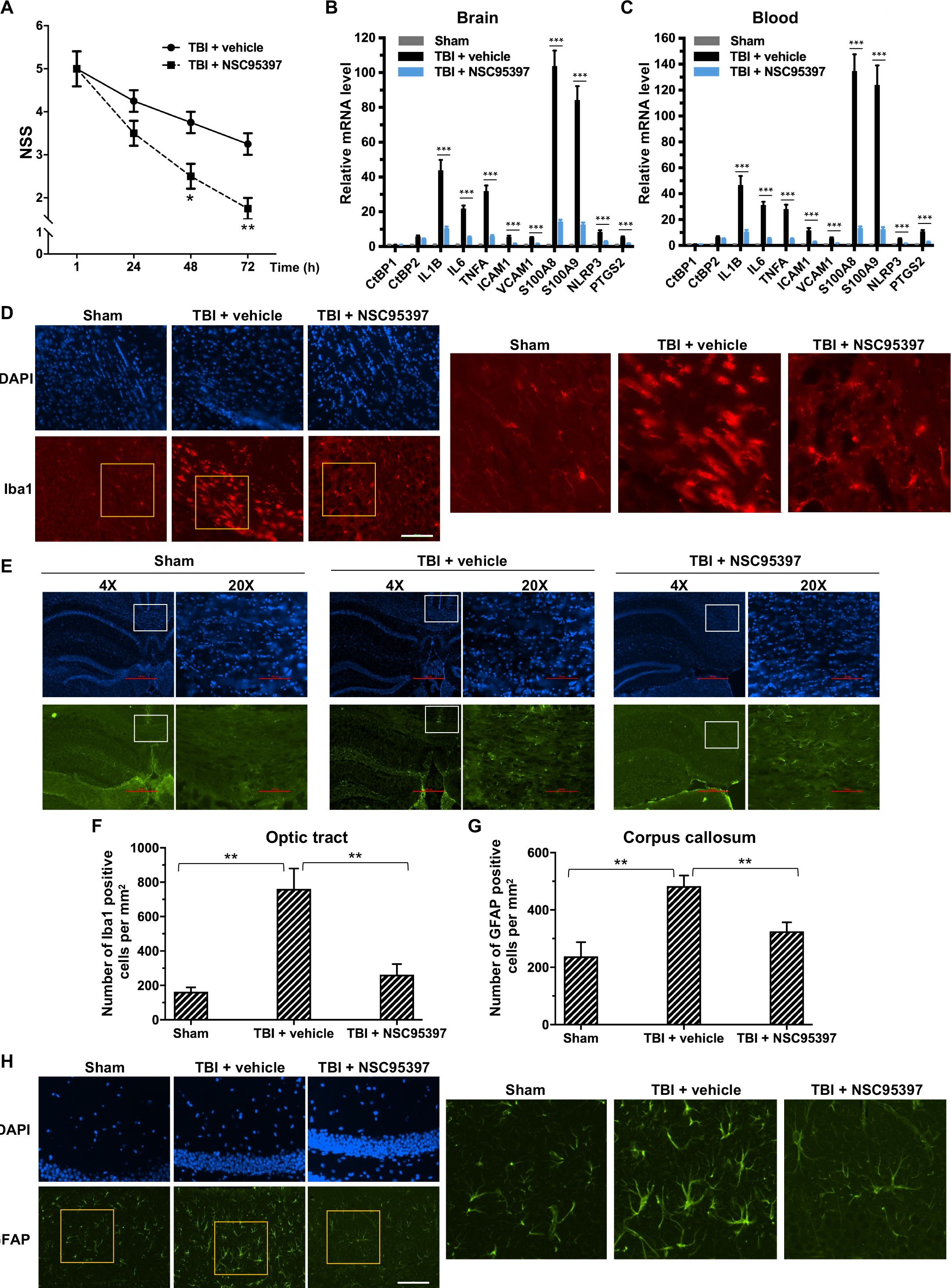
Post-injury treatment with NSC93597 reduces microglia and astrocyte activation, suppresses transactivation of CtBP target genes and improves neurological outcome. (**A**) Comparison of NSS among animals of the sham, TBI and TBI + NSC95397 groups at 1 h, 24 h, 48 h and 72 h postinjury. *n* = 5; \**p*<0.05, \*\**p*<0.01. Mice received i.p. injections of NSC95397 (0.5 mg/kg) at 1 h, 24 h and 48 h after injury. (**B** and **C**) Relative mRNA expression in brain (**B**) and peripheral blood leukocytes (**C**) of the three groups at 72 h postinjury. *n* = 5; \*\*\**p*<0.001. (**D**) Microglial response in the optic tract after single head injury and NSC95397 treatment. Microglia were assessed using Iba1 immunostaining (red) in mouse brain sections prepared near 72 h postinjury. Nuclei were visualized by DAPI staining. Scale bar, 100 μm. The three panels on the right are higher magnification images of the respective framed regions in the Iba1-stained panels on the left. (**E**) Astrocyte response in the corpus callosum visualized by immunostaining for GFAP (green) as described in (**D**). (**F–G**) Quantitation of the microglia and astrocyte response by counting the number of Iba1-positive (**F**) and GFAP-positive (**G**) cells per mm^2^ in the optic tract and corpus callosum regions, respectively. n = 3; \*\**p*<0.01, \*\*\**p*<0.001. (**H**) Comparison of GFAP-positive astrocytes in the hippocampal CA1 region among the three experimental groups. Scale bar, 100 μm. Higher magnification images of the respective framed regions in the GFAP-stained panels on the left are shown in panels on the right.

We next examined the effects of NSC95397 on the response of microglia and astrocytes to local brain injury at 3 d after mTBI by immunofluorescence staining using antibodies specific for the microglial marker Iba1 and the astrocytic marker GFAP (54, 55). Compared to sham brains, injured brains showed significant increases in the number of Iba1-positive microglia in the optic tract and of GFAP-positive astrocytes in the corpus callosum, indicating the activation and proliferation of these CNS glial cells following head injury (Figure 5 D–G). By contrast, we observed significantly reduced numbers of Iba1-positive microglia and GFAP-positive astrocytes in the two above white matter-rich regions of NSC95397-treated brains (Figure 5 D–G). On the other hand, Iba1-positive microglia in the optic tract of the injured brain exhibited the amoeboid-like morphology that is typically associated with activated microglia, while Iba1-positive microglia in the NSC95397-treated brain showed decreased cell soma volume and increased cell ramification, largely resembling the resting state morphology in the sham brain (Figure 5D). In addition to white matter-rich regions, we also noted that NSC95397 treatment features GFAP-positive astrocytes in the hippocampal CA1 region with smaller, more compacted cell bodies and elaborated thinner processes as compared to the vehicle control group (Figure 5H). These results support our hypothesis that the pharmacological inhibition of CtBP1 and CtBP2 is a promising strategy for the treatment of brain injury-induced inflammation and neurodegeneration.

As noted above, another small-molecule CtBP dehydrogenase inhibitor, MTOB, has been shown to antagonize the transcriptional regulatory activity of CtBP1 and CtBP2 by eviction from their target promoters in breast cancer cell lines (49). We therefore investigated whether MTOB, as well as NSC95397, could alleviate neuroinflammation caused by repetitive mild head injury. Mice were given either a single injury (1×TBI) or two injuries 24 h apart (2×TBI). The 2×TBI mice received an injection of placebo (vehicle), MTOB (860 mg/kg) or NSC95397 (1.5 mg/kg) at 1 h and 18 h after the first injury (see Figure 6A for experimental timeline). All mTBI groups showed significantly increased loss of righting reflex (LRR) duration following the first injury as compared to sham mice (Figure 6B). The 1xTBI group regained righting reflex within 24 h after injury, and the 2xTBI group had a significantly longer LRR duration following the second injury (*p* < 0.01) (Figure 6B). The increased duration of righting reflex was effectively suppressed by administration of MTOB (*p* < 0.01) or NSC95397 (*p* < 0.05) after the first injury (Figure 6B). Furthermore, the 2xTBI group exhibited significantly higher NSS than the 1xTBI group at 24 h, 48 h and 72 h (*p* < 0.001) following the first injury (Figure 6C). NSS scores were significantly decreased by MTOB (*p* < 0.001 at 24 h; *p* < 0.01 at 48 h; and *p* < 0.05 at 72 h) and by NSC95397 (*p* < 0.001 at 24 h and *p* < 0.05 at 48 h) (Figure 6C). In addition, we observed a 150 to 230% increase in the mRNA expression of the nine CtBP-regulated proinflammatory genes in the brain tissues of the 2xTBI group as compared to the 1xTBI group at 3 d after the first injury (Figure 6D). The inhibitor-treated mice exhibited lower or equal levels of expression of these CtBP target genes relative to the 1xTBI mice (Figure 6D). Therefore, MTOB and NSC95397 attenuate repetitive head injury-elicited neurologic dysfunction via inhibition of the transactivation activity of CtBP1 and CtBP2.

**Figure 6.**
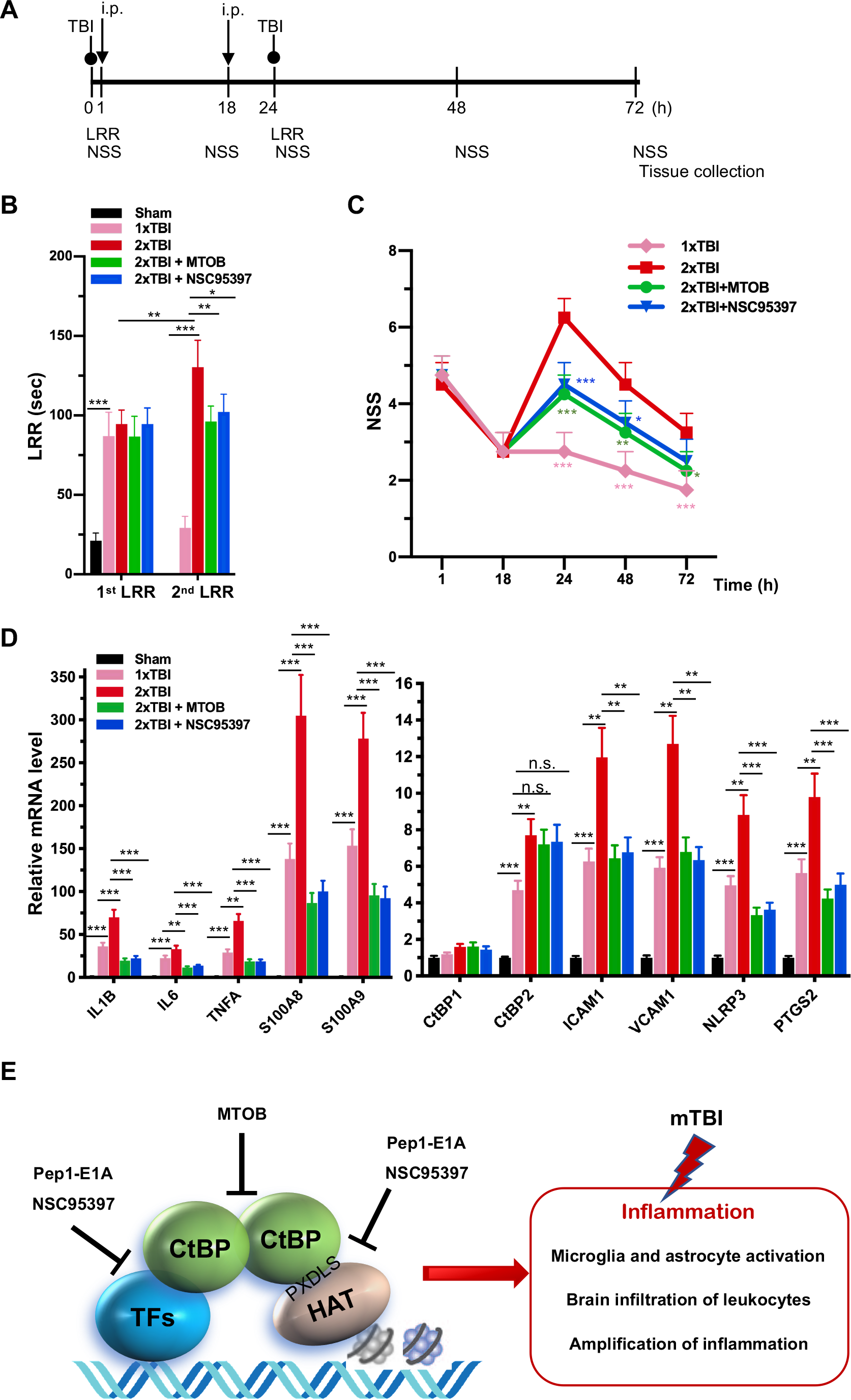
NSC95397 and MTOB attenuate neuroinflammation caused by repetitive mild TBI. (**A**) Experimental timeline. Mice received a single head impact of 0.5 J energy (1xTBI), or two 0.5 J impacts (2xTBI) spaced 24 h apart. The CtBP inhibitor-treated groups were given an i.p. injection of MTOB (860 mg/kg) or NSC95397 (1.5 mg/kg) at 1 h and 18 h after the first injury. (**B**) Comparison of LRR durations following the first and second head injury. *n* = 5; \**p*<0.05, \*\**p*<0.01. \*\*\**p*<0.001. (**C**) CtBP inhibitors improved neurological deficits in mice receiving repeated mTBI. NSS assessment at 1 h and 18 h were given prior to the administration of the CtBP inhibitors. *n* = 5; \**p*<0.05, \*\**p*<0.01, \*\*\**p*<0.001. (**D**) NSC95397 and MTOB prevent a further increase in the mRNA expression levels of CtBP target genes in the animal brains with repeated TBI. Brain tissues were collected for mRNA analysis at 72 h after the first injury; results were normalized to sham. *n* = 5; \*\**p*<0.01, \*\*\**p*<0.001. (**E**) A model for the transcriptional activation and its inhibition involving the CtBPs, HAT and transcription factors (TFs) in TBI neuroimmunology. CtBP-mediated, TBI-dependent proinflammatory gene induction necessitates CtBP activation through dimerization.

In summary, we prove that CtBP1 and CtBP2 can transactivate select proinflammatory target genes in single and repetitive mTBI mice, thus contributing to acute neurological deficits resembling clinical mTBI symptoms that can be quantified using the NSS score, as in our model (see Figure 6E). We have also demonstrated that acute treatment with CtBP inhibitors mitigates the TBI-induced inflammatory pathology and significantly improves mouse brain function. Finally, and most importantly, these findings suggest that suppression of CtBP function could potentially be an effective strategy to alleviate TBI-induced chronic inflammation and neurological impairment.

## Materials and Methods

### Animals

All animal experiments were reviewed and approved by the University of Colorado Anschutz Medical Campus Institutional Animal Care and Use Committee (protocol B-114116(12)1E) and performed in accordance with relevant guidelines and regulations. C57BL/6 mice were group housed in environmentally controlled conditions with 12:12 h light:dark cycle and provided food and water *ad libitum*.

### CHIMERA TBI

Mice at 16–20 weeks old of age (body weight 20–30 g) were randomly assigned to sham, injury, or injury with treatment groups. Animals were anaesthetized with isoflurane (induction: 4.5%, maintenance: 2.5-3%) in oxygen (0.9 L/min) and underwent TBI or sham procedures as described (50). The average total duration of isoflurane exposure for the whole procedure was 3 to 4 min. For single TBI experiments, mice received a closed-head impact from a pneumatically driven 50 g steel piston that was calibrated to deliver 0.5 to 0.8 J of kinetic energy at the point of impact. An impact of 0.5 J energy was defined as the low threshold for producing CHIMERA-delivered mild brain injury in a previous dose-response study (56). In the repeated mTBI experiment, mice received two successive 0.5 J impacts spaced 24 h apart. The sham group underwent all procedures except for the impact. For receiving head injury, the animal was placed in a supine position in the holding bed with the top of its head placed flat over a hole in the head plate. The top of the animal’s head was aligned using crosshairs so that the piston strikes a circular area with a radius of 2.5 mm surrounding the bregma on the vertex of the head. In the control experiment to distinguish between the local and systemic effect of CHIMERA-delivered impact (see Figure S2), animals were held in place such that the animal’s back or one ear was positioned over the hole of the head plate to receive the piston-delivered impact.

### Administration of CtBP inhibitors

Preparation of the PLDLS-containing Pep1-E1A and the PLDLS-to-PLDEL mutant (Pep1-E1A^Mut^) peptides were as previously described (41). Peptides (6 mg/mL) and MTOB sodium salt (Sigma, K6000; 425 mg/mL) were dissolved in water. NSC95397 (Sigma, N1786; 5 mg/mL) was dissolved in DMSO. The stock solutions were diluted in PBS solution (vehicle) to desired final concentrations freshly before use. Pep1-E1A was delivered at 2 mg/kg dosage, 4-methylthio-2-oxobutyric acid (MTOB) sodium salt at 860 mg/kg and NSC95397 at 0.75 or 1.5 mg/kg with intraperitoneal (i.p.) injection.

### Behavioral analyses

The latency to righting reflex (a.k.a. loss of righting reflex, LRR), a surrogate measurement for loss of consciousness after brain injury, is used as a behavioral indicator of injury severity (57). The LRR duration was recorded as the time interval from the discontinuation of isoflurane exposure until the animal spontaneously rights itself from a supine position (50). Post-traumatic neurological impairments were evaluated using a 10-point neurological severity score (NSS) paradigm, which evaluate mice in performance of ten tasks: circle exiting, seeking behavior, gait pattern (monoparesis or hemiparesis) and paw grip, straight walk, startle reflex, beam (7 mm × 7 mm) balancing, beam walking on a flat surface of 3, 2 and 1 cm width, and round stick (5 mm in diameter) balancing (58). One point is given for the lack of a tested reflex or the inability to perform the task, with a maximal NSS of 10 points indicating the failure of all tasks.

### Tissue collection, histology, and immunofluorescence

Mice were anesthetized with isoflurane via a nose cone. Blood samples were collected by cardiac puncture into EDTA-coated tube to prevent clotting. Leukocytes were collected by two rounds of 10-min incubation in a red blood cells lysis buffer (0.8% NH_4_Cl, 0.084% NaHCO_3_ and 0.037% EDTA) followed by centrifugation at 2500 × g for 10 min, all at room temperature. The pelleted white blood cells were frozen in liquid nitrogen and stored at −80°C until RNA extraction. The animals were transcardially perfused with ice-cold 0.8% NaCl solution for 4 to 5 min until the liver is cleared of blood. Mouse brain was obtained by decapitation of the perfused animal. For biochemical analyses, the brain was longitudinally hemisected, rapidly frozen in liquid nitrogen and stored at −80°C until RNA and protein extraction. For immunohistological analyses, mice were perfused with PBS followed by 4% ice-cold paraformaldehyde-PBS solution for 5 min. Brain tissue were post-fixed in 10% formalin-PBS for 48 h at 4°C, embedded in paraffin, and sectioned coronally to 40-μm thickness. The primary antibodies used for immunofluorescence imaging were: mouse anti-CtBP2 (1:200,BD, 612044), goat anti-CtBP2 (Santa Cruz, sc-5966, 1:100), rabbit anti-Iba1(GeneTex, GTX100042, 1:200), rabbit anti-GFAP (Abcam, AB7260, 1:1000), mouse anti-NeuN (GeneTex, GTX30773, 1:100). Secondary antibodies used were IgG conjugated to Alexa Flour 594 or 488 (1:500, Invitrogen), biotin-labeled IgG (1:300, Vector lab), Streptavidin conjugated to Alexa Flour 594 or 488 (1:2000, Life Technology, S11227 and S11223). Mouse on Mouse (MOM) Detection kit (Vector Laboratories, BMK-2202) were used to reduce endogenous mouse Ig straining when using mouse primary antibodies on mouse tissues. All slides were counterstained with DAPI to visualize the nuclei.

### Cell transfection and LPS treatment

The mouse macrophage cell line RAW264.7 was obtained from American Type Culture Collection (ATCC, VA, USA). The mouse microglial BV2 cell line was a kind gift from Dr. Manisha Patel (University of Colorado Anschutz Medical Campus). Both cell lines were cultured in Dulbecco's modified Eagle’s medium (DMEM; Corning 10017CV) supplemented with 10% fetal bovine serum (FBS, GEMINI, 100-500) and 1% penicillin/streptomycin (Corning, 30002CI) at 37°C with 5% CO_2_.

Cell transfection was carried out in 24-well plates as described (59). Briefly, RAW264.7 and BV2 cells under approximately 80% confluence were treated with 0.25% Trypsin-EDTA (Corning, 25-053-CI) and then transfected with Lipofectamine 2000 (Invitrogen, 11668019) in suspension with siRNAs specific to mouse *CtBP1* (UCUUCCACAGUGUGACUGCGUUAUUUU, 50 nM), *CtBP2* (GCCUUUGGAUUCAGCGUCAUAUUU, 50 nM), or both (25 nM each). The transfected cells were incubated in DMEM for 24 h, followed by treatment with 200 ng/mL LPS (Sigma-Aldrich, L3880) for 6 h, and harvested for RNA extraction.

### Primary microglia and astrocyte cultures and treatment with peptide and LPS

Mouse primary microglia and astrocytes were isolated from cerebral cortices of neonatal pups (P0 to P2-d-old) and cultured as described (60, 61), with minor modifications. The isolated cortices were washed with ice-cold PBS containing 0.1% BSA and 1 × minimum essential medium (MEM) non-essential amino acids (Corning, 25025CL). Single cell suspension was made by pipetting up and down through a sterile 10 ml pipette ten times followed by passing through a 18G needle three times. The mixed cortical cells were passed through a nylon mesh cell strainer with 70-μm pores and plated on an uncoated T75 flask in DMEM (Corning, 10013CV) with 10% FBS (Gemini, 100-500). After the mixed glial culture reached confluency at 10 to 14 d post-plating, microglia were detached using an orbital shaker (220 rpm, 1 h), collected and re-plated in DMEM with 15% FBS. Oligodendrocyte precursor cells were removed by further shaking at 220 rpm for 6 h. The remaining astrocyte layer were detached by trypsin digestion and re-plated in DMEM with 10% FBS. The primary microglia and astrocyte cultures were >90% pure based on immunofluorescence imaging with antibodies specific to Iba1 and GFAP, respectively.

Cultured primary microglia or astrocytes were incubated with 20 μM Pep1-E1A or Pep1-E1A^Mut^ for 2 h before the addition of 200 ng/mL LPS (Sigma-Aldrich, L3880). Cells were harvested 2 h after LPS treatment for RNA extraction. Anti-FLAG immunofluorescence staining was performed to monitor cellular internalization of the peptides as described (41).

### Reverse Transcription and Quantitative real-time PCR (RT-qPCR)

Total RNA was extracted from freshly harvested cells or frozen tissues using TRIzol^TM^ reagent (Invitrogen, 15596026) and analyzed by RT-qPCR. First-strand cDNA was reverse transcribed from 1.0 μg total RNA with oligo (dT) primers using Verso cDNA synthesis kit (Thermo Scientific, AB1453A). Quantitative PCR with SYBR green detection (Applied Biosystems, A25741) was performed using 1% of the reversely transcribed cDNA mixture on a BioRad CFX96 real-time detection system. Relative expression of individual genes were normalized to *ACTB* (β-Actin) expression using the 2^−ΔΔCt^ method (62). The sequences of all RT-qPCR primers were provided in Supplementary Table 1.

### Chromatin immunoprecipitation (ChIP)

ChIP experiments were carried out as described (63) with modifications. In brief, cells were cross-linked with 1% (w/v) formaldehyde for 15 min at room temperature. The cross-linking was quenched with 0.125 M glycine for 5 min. About 1 × 10^8^ cross-linked cells were lysed by sonication for 15 × 30 s with 30 s break on ice in 10 mL lysis buffer (0.1% SDS, 0.5% Triton X-100, 150 mM NaCl, 20 mM Tris-pH 8.1, 1 mM DTT, 2 mM EDTA, and with protease inhibitor cocktail freshly added before use; Roche, 11836153001). After centrifugation at 19,000 × g for 15 min, the supernatant was used for immunoprecipitation with 1/100 of the lysate (100 μL) set aside as the input sample. The remaining lysate were split into four equal parts and incubated with each antibody (mouse anti-CtBP1, BD Biosciences, 612042; mouse anti-CtBP2, BD Biosciences, 612044; mouse anti-p300 Santa Cruz, sc-48343; and mouse IgG_1_, Santa Cruz, sc-69786) overnight with rotation at 4°C. Protein A beads (RepliGen, 10-1003-03) were added to the lysate with rotation at 4°C for 4 h. The beads were washed 5 × 5 min with the lysis buffer. The precipitated DNA-protein complex was eluted into 300 μL 1 mM NaHCO_3_ and 1% SDS by incubation at 65°C for 15 min. After centrifugation at 19,000 × g for 1 min, the supernatant was transferred to fresh tubes, followed by supplementation with 5 M NaCl to a final concentration of 300 mM and incubation at 65°C overnight to reverse the cross-linking. Protein and SDS of both input and IP samples were removed through phenol-chloroform extraction and ethanol precipitation. The DNA of input and ChIP samples was resuspended in 100 μL and 400 μL H_2_O, respectively. For qPCR, 1.5 μL from each sample was used. For each antibody, relative ChIP signal was calculated as percent of input using the 2^−ΔΔCt^ method (62) and the ChIP signal of the LPS-stimulated cells were normalized to that of the non-stimulated control. The sequences of all ChIP-qPCR primers were provided in Supplemental Table 1.

### Western Blotting

Protein extracts were prepared from longitudinally halved mouse brain (~200 mg) in 500 μL homogenization buffer consisting of 10 mM Tris, pH 7.4, 100 mM NaCl, 1 mM EDTA, 1 mM EGTA, 1% Triton X-100, 10% glycerol, 0.1% SDS, 0.5% deoxycholate, and 1 × complete protease inhibitor cocktail (Roche, 11836153001). Debris were removed by centrifugation at 18,000 × g for 5 min at 4°C. The supernatant fraction was quantified through use of the Bradford assay (Bio-Rad, 5000006). For each sample, 50 μg of total protein extracts was resolved by 12% SDS-PAGE and transferred onto a PVDF membrane for immunodetection. Membrane were incubated with antibodies specific to CtBP1 (BD Biosciences, 612042, 1:1,000), CtBP2 (BD Biosciences, 612044, 1:1,000), S100A9 (Santa Cruz Biotechnology, sc-58706, 1:1,000), NLRP3 (Santa Cruz Biotechnology, 134306, 1:1,000) and GAPDH (Sigma, G8795, 1:5,000) at 4°C overnight, followed by peroxidase-labeled appropriate secondary antibodies at room temperature for 1 h. The membrane was developed using an enhanced chemiluminescence substrate (Millipore Corporation, WBKLS0500) and scanned with a ChemiDoc MP imager (Bio-Rad). Raw signal intensity for each band was measured using Image J software (4.0.1 version).

### Statistical Analysis

Plots were made and statistical analysis was performed using GraphPad Prism version 8.0 (GraphPad Software). Data are expressed as mean ± standard deviation (SD). In cell culture experiments the ‘*n*’ values denote the number of experiments. Each independent experiment contained triplicate cultured wells.

## Acknowledgements

We thank H. Potter, Y. Shellman, X-J. Wang and D. Xue for comments on the manuscript. This work was supported by the Dudley Stem Cell Research Fund of the University of Colorado Cancer Center to M. H. and a research grant from the VA Eastern Colorado Health Care System (5101BX002370) to D.N. R. Z was supported in part by NIH grants R01CA221282, R01GM126157, and R01GM114178. M.B. was supported by the Lung, Head and Neck Cancer training grant T32CA174648 and an American Lung Association Senior Research Training Fellowship.

---

**Figure S1.**
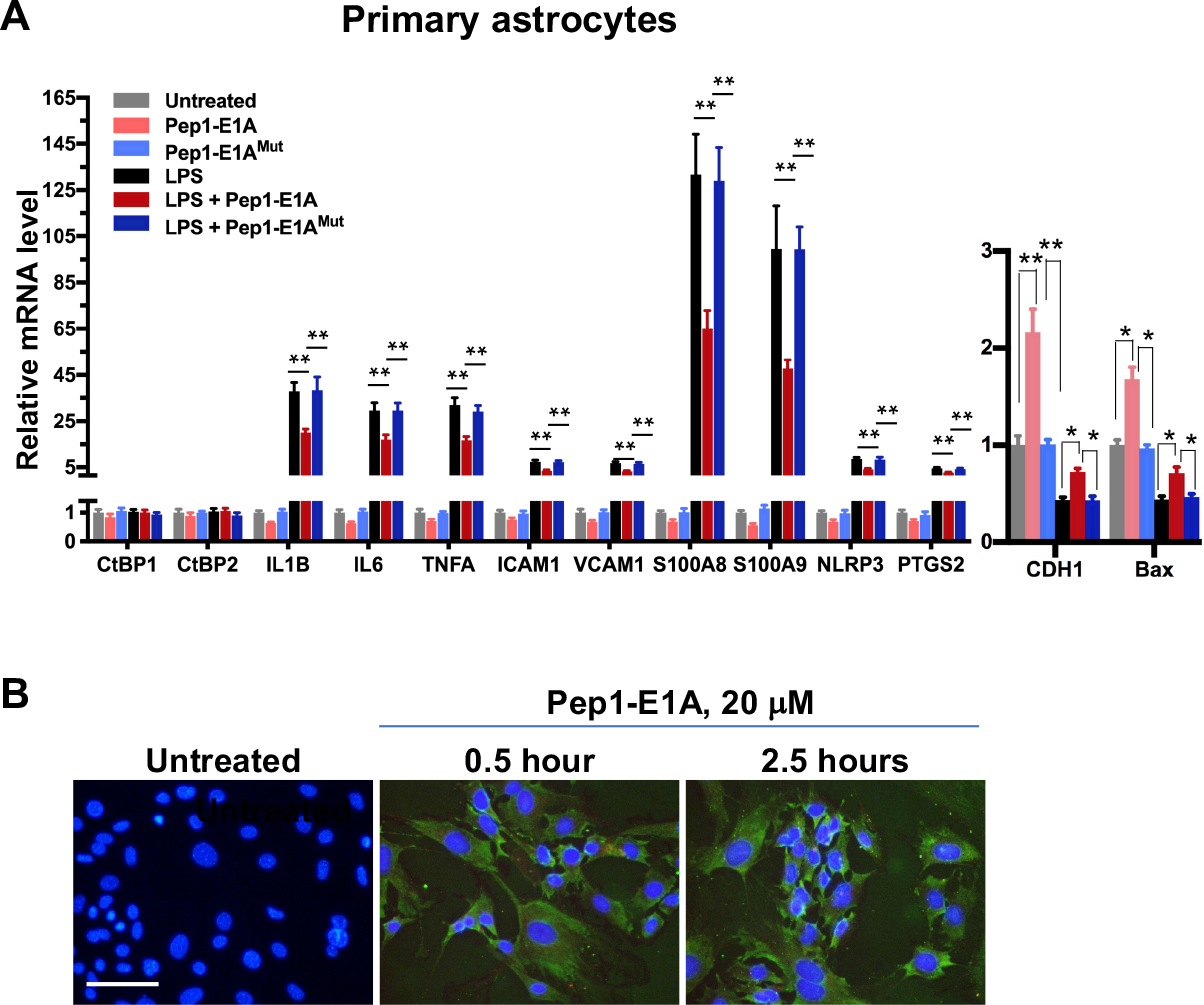
Pep1-E1A suppresses LPS-induced inflammatory gene expression in mouse primary astrocytes. (**A**) Effects of pretreatment with 20 μM Pep1-E1A or Pep1-E1A^Mut^ on the basal and LPS-induced mRNA expression of CtBP target genes in mouse primary astrocytes. *n* = 3; \**p*<0.05, \*\**p*<0.01, \*\*\**p*<0.001. (**B**) Representative immunostaining images of Pep1-E1A internalization into cultured astrocytes after 0.5 and 2.5 h incubation. Scale bar, 50 μm.

**Figure S2.**
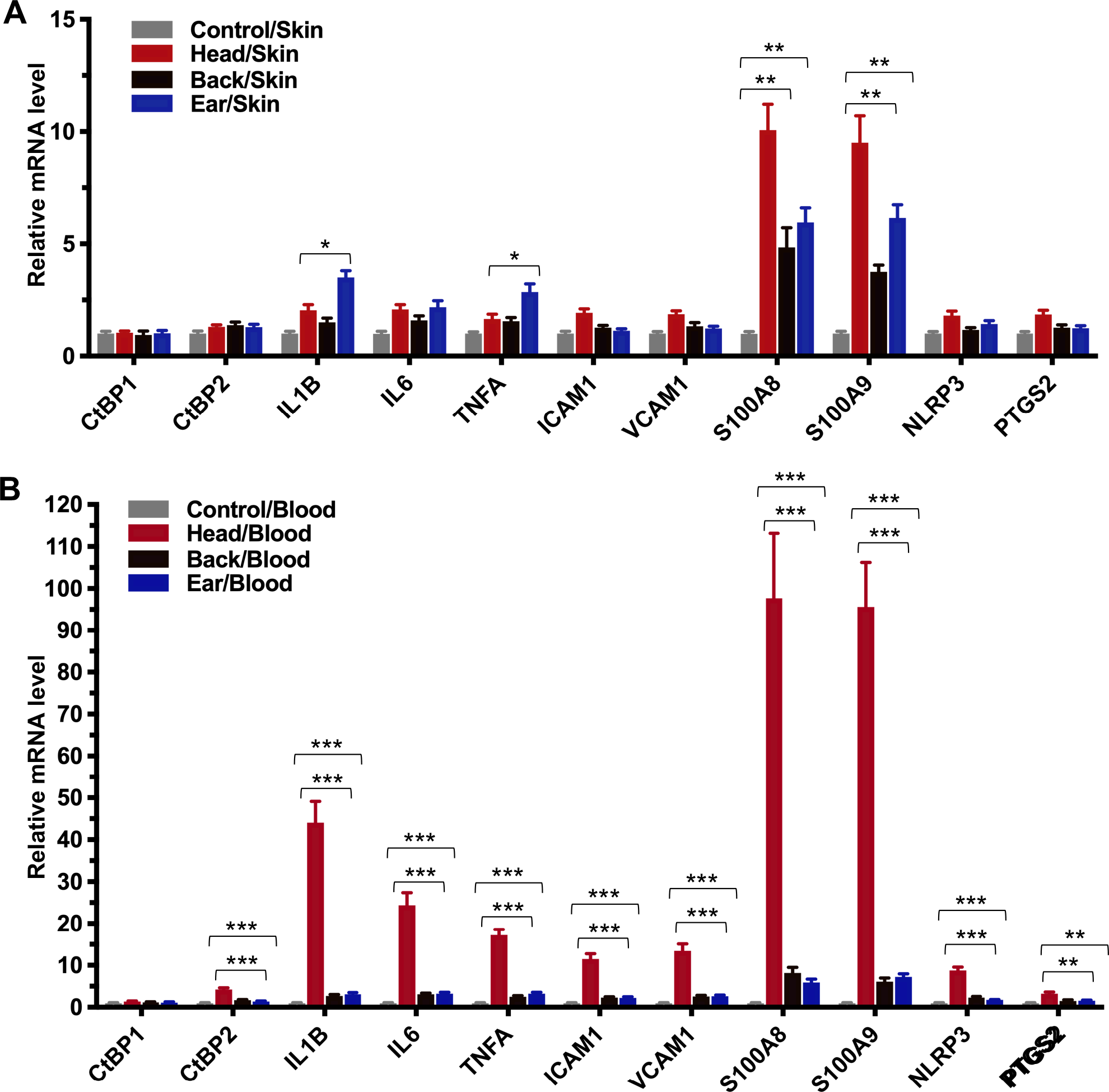
CHIMERA-delivered mild TBI causes systemic inflammatory response. Mice received sham procedure (control) or a single impact of 0.7 J energy to the head, the back, or the ear. Skin tissues containing the site of the impact (**A**) and peripheral blood leukocytes (**B**) and were collected 24 h postinjury for total RNA extraction and RT-qPCR analysis. *n* = 4; \**p*<0.05, \*\**p*<0.01, \*\*\**p*<0.001.

## References

1. Faul M & Coronado V (2015) Epidemiology of traumatic brain injury. Handb Clin Neurol 127:3–13.

2. Smith DH, Johnson VE, & Stewart W (2013) Chronic neuropathologies of single and repetitive TBI: substrates of dementia? Nat Rev Neurol 9(4):211–221.

3. Gardner RC & Yaffe K (2014) Traumatic brain injury may increase risk of young onset dementia. Ann Neurol 75(3):339–341.

4. Bramlett HM & Dietrich WD (2015) Long-Term Consequences of Traumatic Brain Injury: Current Status of Potential Mechanisms of Injury and Neurological Outcomes. J Neurotrauma 32(23):1834–1848.

5. Gardner RC & Yaffe K (2015) Epidemiology of mild traumatic brain injury and neurodegenerative disease. Mol Cell Neurosci 66(Pt B):75–80.

6. Godbolt AK, et al. (2014) Systematic review of the risk of dementia and chronic cognitive impairment after mild traumatic brain injury: results of the International Collaboration on Mild Traumatic Brain Injury Prognosis. Arch Phys Med Rehabil 95(3 Suppl):S245–256.

7. Boyle E, et al. (2014) Systematic review of prognosis after mild traumatic brain injury in the military: results of the International Collaboration on Mild Traumatic Brain Injury Prognosis. Arch Phys Med Rehabil 95(3 Suppl):S230–237.

8. Cancelliere C, et al. (2014) Systematic review of prognosis and return to play after sport concussion: results of the International Collaboration on Mild Traumatic Brain Injury Prognosis. Arch Phys Med Rehabil 95(3 Suppl):S210–229.

9. Carroll LJ, et al. (2014) Systematic review of the prognosis after mild traumatic brain injury in adults: cognitive, psychiatric, and mortality outcomes: results of the International Collaboration on Mild Traumatic Brain Injury Prognosis. Arch Phys Med Rehabil 95(3 Suppl):S152–173.

10. McCauley SR, et al. (2014) Patterns of early emotional and neuropsychological sequelae after mild traumatic brain injury. J Neurotrauma 31(10):914–925.

11. Mittl RL, et al. (1994) Prevalence of MR evidence of diffuse axonal injury in patients with mild head injury and normal head CT findings. AJNR Am J Neuroradiol 15(8):1583–1589.

12. Topal NB, et al. (2008) MR imaging in the detection of diffuse axonal injury with mild traumatic brain injury. Neurol Res 30(9):974–978.

13. Steenkamp MM, Litz BT, Hoge CW, & Marmar CR (2015) Psychotherapy for Military-Related PTSD: A Review of Randomized Clinical Trials. JAMA 314(5):489–500.

14. Barnes DE, et al. (2018) Association of Mild Traumatic Brain Injury With and Without Loss of Consciousness With Dementia in US Military Veterans. JAMA Neurol 75(9):1055–1061.

15. McKee AC, Daneshvar DH, Alvarez VE, & Stein TD (2014) The neuropathology of sport. Acta Neuropathol 127(1):29–51.

16. Simon DW, et al. (2017) The far-reaching scope of neuroinflammation after traumatic brain injury. Nat Rev Neurol 13(3):171–191.

17. Russo MV & McGavern DB (2016) Inflammatory neuroprotection following traumatic brain injury. Science 353(6301):783–785.

18. Jassam YN, Izzy S, Whalen M, McGavern DB, & El Khoury J (2017) Neuroimmunology of Traumatic Brain Injury: Time for a Paradigm Shift. Neuron 95(6):1246–1265.

19. Narayana PA (2017) White matter changes in patients with mild traumatic brain injury: MRI perspective. Concussion 2(2):CNC35.

20. Fehily B & Fitzgerald M (2017) Repeated Mild Traumatic Brain Injury: Potential Mechanisms of Damage. Cell Transplant 26(7):1131–1155.

21. Witcher KG, Eiferman DS, & Godbout JP (2015) Priming the inflammatory pump of the CNS after traumatic brain injury. Trends Neurosci 38(10):609–620.

22. Gruenbaum SE, Zlotnik A, Gruenbaum BF, Hersey D, & Bilotta F (2016) Pharmacologic Neuroprotection for Functional Outcomes After Traumatic Brain Injury: A Systematic Review of the Clinical Literature. CNS Drugs 30(9):791–806.

23. McConeghy KW, Hatton J, Hughes L, & Cook AM (2012) A review of neuroprotection pharmacology and therapies in patients with acute traumatic brain injury. CNS Drugs 26(7):613–636.

24. Jayakumar AR, et al. (2014) Activation of NF-kappaB mediates astrocyte swelling and brain edema in traumatic brain injury. J Neurotrauma 31(14):1249–1257.

25. Mettang M, et al. (2018) IKK2/NF-kappaB signaling protects neurons after traumatic brain injury. FASEB J 32(4):1916–1932.

26. Bergman LM & Blaydes JP (2006) C-terminal binding proteins: emerging roles in cell survival and tumorigenesis. Apoptosis 11(6):879–888.

27. Chinnadurai G (2009) The transcriptional corepressor CtBP: a foe of multiple tumor suppressors. Cancer research 69(3):731–734.

28. Dcona MM, Morris BL, Ellis KC, & Grossman SR (2017) CtBP- an emerging oncogene and novel small molecule drug target: Advances in the understanding of its oncogenic action and identification of therapeutic inhibitors. Cancer Biol Ther 18(6):379–391.

29. Quinlan KG, et al. (2006) Specific recognition of ZNF217 and other zinc finger proteins at a surface groove of C-terminal binding proteins. Molecular and cellular biology 26(21):8159–8172.

30. Nardini M, et al. (2003) CtBP/BARS: a dual-function protein involved in transcription co-repression and Golgi membrane fission. The EMBO journal 22(12):3122–3130.

31. Bruton RK, et al. (2008) Identification of a second CtBP binding site in adenovirus type 5 E1A conserved region 3. J Virol 82(17):8476–8486.

32. Kim JH, Cho EJ, Kim ST, & Youn HD (2005) CtBP represses p300-mediated transcriptional activation by direct association with its bromodomain. Nat Struct Mol Biol 12(5):423–428.

33. Kuppuswamy M, et al. (2008) Role of the PLDLS-binding cleft region of CtBP1 in recruitment of core and auxiliary components of the corepressor complex. Molecular and cellular biology 28(1):269–281.

34. Ray SK, Li HJ, Metzger E, Schule R, & Leiter AB (2014) CtBP and associated LSD1 are required for transcriptional activation by NeuroD1 in gastrointestinal endocrine cells. Molecular and cellular biology 34(12):2308–2317.

35. Shi Y, et al. (2003) Coordinated histone modifications mediated by a CtBP co-repressor complex. Nature 422(6933):735–738.

36. Balasubramanian P, Zhao LJ, & Chinnadurai G (2003) Nicotinamide adenine dinucleotide stimulates oligomerization, interaction with adenovirus E1A and an intrinsic dehydrogenase activity of CtBP. FEBS letters 537(1-3):157–160.

37. Madison DL, Wirz JA, Siess D, & Lundblad JR (2013) Nicotinamide adenine dinucleotide-induced multimerization of the co-repressor CtBP1 relies on a switching tryptophan. The Journal of biological chemistry 288(39):27836–27848.

38. Schaeper U, et al. (1995) Molecular cloning and characterization of a cellular phosphoprotein that interacts with a conserved C-terminal domain of adenovirus E1A involved in negative modulation of oncogenic transformation. Proceedings of the National Academy of Sciences of the United States of America 92(23):10467–10471.

39. Blevins MA, Huang M, & Zhao R (2017) The Role of CtBP1 in Oncogenic Processes and Its Potential as a Therapeutic Target. Mol Cancer Ther 16(6):981–990.

40. Byun JS & Gardner K (2013) C-Terminal Binding Protein: A Molecular Link between Metabolic Imbalance and Epigenetic Regulation in Breast Cancer. International journal of cell biology 2013:647975.

41. Blevins MA, et al. (2018) CPP-E1A fusion peptides inhibit CtBP-mediated transcriptional repression. Mol Oncol 12(8):1358–1373.

42. Patel J, et al. (2014) Inhibition of C-terminal binding protein attenuates transcription factor 4 signaling to selectively target colon cancer stem cells. Cell cycle 13(22):3506–3518.

43. Straza MW, et al. (2010) Therapeutic targeting of C-terminal binding protein in human cancer. Cell cycle 9(18):3740–3750.

44. Achouri Y, Noel G, & Van Schaftingen E (2007) 2-Keto-4-methylthiobutyrate, an intermediate in the methionine salvage pathway, is a good substrate for CtBP1. Biochem Biophys Res Commun 352(4):903–906.

45. Boxer LD, Barajas B, Tao S, Zhang J, & Khavari PA (2014) ZNF750 interacts with KLF4 and RCOR1, KDM1A, and CTBP1/2 chromatin regulators to repress epidermal progenitor genes and induce differentiation genes. Genes & development 28(18):2013–2026.

46. Paliwal S, Ho N, Parker D, & Grossman SR (2012) CtBP2 Promotes Human Cancer Cell Migration by Transcriptional Activation of Tiam1. Genes Cancer 3(7-8):481–490.

47. Ashburner M, et al. (2000) Gene ontology: tool for the unification of biology. The Gene Ontology Consortium. Nat Genet 25(1):25–29.

48. The Gene Ontology C (2019) The Gene Ontology Resource: 20 years and still GOing strong. Nucleic acids research 47(D1):D330–D338.

49. Di LJ, et al. (2013) Genome-wide profiles of CtBP link metabolism with genome stability and epithelial reprogramming in breast cancer. Nature communications 4:1449.

50. Namjoshi DR, et al. (2014) Merging pathology with biomechanics using CHIMERA (Closed-Head Impact Model of Engineered Rotational Acceleration): a novel, surgery-free model of traumatic brain injury. Mol Neurodegener 9:55.

51. Csuka E, et al. (1999) IL-10 levels in cerebrospinal fluid and serum of patients with severe traumatic brain injury: relationship to IL-6, TNF-alpha, TGF-beta1 and blood-brain barrier function. J Neuroimmunol 101(2):211–221.

52. Helmy A, Carpenter KL, Menon DK, Pickard JD, & Hutchinson PJ (2011) The cytokine response to human traumatic brain injury: temporal profiles and evidence for cerebral parenchymal production. J Cereb Blood Flow Metab 31(2):658–670.

53. Vourc’h M, Roquilly A, & Asehnoune K (2018) Trauma-Induced Damage-Associated Molecular Patterns-Mediated Remote Organ Injury and Immunosuppression in the Acutely Ill Patient. Front Immunol 9:1330.

54. Bignami A & Dahl D (1974) Astrocyte-specific protein and neuroglial differentiation. An immunofluorescence study with antibodies to the glial fibrillary acidic protein. J Comp Neurol 153(1):27–38.

55. Ito D, et al. (1998) Microglia-specific localisation of a novel calcium binding protein, Iba1. Brain Res Mol Brain Res 57(1):1–9.

56. Namjoshi DR, et al. (2017) Defining the biomechanical and biological threshold of murine mild traumatic brain injury using CHIMERA (Closed Head Impact Model of Engineered Rotational Acceleration). Exp Neurol 292:80–91.

57. Dewitt DS, Perez-Polo R, Hulsebosch CE, Dash PK, & Robertson CS (2013) Challenges in the development of rodent models of mild traumatic brain injury. J Neurotrauma 30(9):688–701.

58. Flierl MA, et al. (2009) Mouse closed head injury model induced by a weight-drop device. Nature protocols 4(9):1328–1337.

59. Dalby B, et al. (2004) Advanced transfection with Lipofectamine 2000 reagent: primary neurons, siRNA, and high-throughput applications. Methods 33(2):95–103.

60. Lian H, Roy E, & Zheng H (2016) Protocol for Primary Microglial Culture Preparation. Bio Protoc 6(21).

61. Schildge S, Bohrer C, Beck K, & Schachtrup C (2013) Isolation and culture of mouse cortical astrocytes. J Vis Exp (71).

62. Winer J, Jung CK, Shackel I, & Williams PM (1999) Development and validation of real-time quantitative reverse transcriptase-polymerase chain reaction for monitoring gene expression in cardiac myocytes in vitro. Anal Biochem 270(1):41–49.

63. Huang M, Zhou Z, & Elledge SJ (1998) The DNA replication and damage checkpoint pathways induce transcription by inhibition of the Crt1 repressor. Cell 94(5):595–605.

